# The chromosomal-level genome assembly and comprehensive transcriptomes of Chinese razor clam (*Sinonovacula constricta*) with deep-burrowing life style and broad-range salinity adaptation

**DOI:** 10.1101/735142

**Authors:** Yinghui Dong, Qifan Zeng, Jianfeng Ren, Hanhan Yao, Wenbin Ruan, Liyuan Lv, Lin He, Qinggang Xue, Zhenmin Bao, Shi Wang, Zhihua Lin

## Abstract

**Background:** The Chinese razor clam, *Sinonovacula constricta*, is one of the commercially important marine bivalves with deep-burrowing lifestyle and remarkable adaptability of broad-range salinity. Despite its economic impact and representative of the less-understood deep-burrowing bivalve lifestyle, there are few genomic resources for exploring its unique biology and adaptive evolution. Herein, we reported a high-quality chromosomal-level reference genome of *S. constricta*, the first genome of the family Solenidae, along with a large amount of short-read/full-length transcriptomic data of whole-ontogeny developmental stages, all major adult tissues, and gill tissues under salinity challenge.

**Findings:** A total of 101.79 Gb and 129.73 Gb sequencing data were obtained with the PacBio and Illumina platforms, which represented approximately 186.63X genome coverage. In addition, a total of 160.90 Gb and 24.55 Gb clean data were also obtained with the Illumina and PacBio platforms for transcriptomic investigation. A *de novo* genome assembly of 1,340.13 Mb was generated, with a contig N50 of 689.18 kb. Hi-C scaffolding resulted in 19 chromosomes with a scaffold N50 of 57.99 Mb. The repeat sequences account for 50.71% of the assembled genome. A total of 26,273 protein-coding genes were predicted and 99.5% of them were annotated. Phylogenetic analysis revealed that *S. constricta* diverged from the lineage of Pteriomorphia at approximately 494 million years ago. Notably, cytoskeletal protein tubulin and motor protein dynein gene families are rapidly expanded in the *S. constricta* genome and are highly expressed in the mantle and gill, implicating potential genomic bases for the well-developed ciliary system in the *S. constricta*.

**Conclusions:** The high-quality genome assembly and comprehensive transcriptomes generated in this work not only provides highly valuable genomic resources for future studies of *S. constricta*, but also lays a solid foundation for further investigation into the adaptive mechanisms of benthic burrowing mollusks.

## Introduction

The Chinese razor clam *Sinonovacula constricta* (Lamarck 1818) is a member of the Solenidae family of bivalve molluscs, recognizing for its straight razor-like shape and fragile shells (Figure 1A). It is widely distributed in the intertidal zone along the west Pacific Ocean and engages in a pelago-benthic life cycle (Figure 1B). As adaptation to a deep-burrowing lifestyle, the razor clam is characterized by smooth shells, muscular foot, and elongated siphons (Figure 1). Benefit from its relatively short production cycle and high productive efficiency, the razor clam has become one of the four most important maricultured bivalve species (together with oyster, scallop, and *Venerupis* spp.) in China, Japan, and Korea, with over 800,000 metric tons of production in 2016 (FAO, 2018).

**Figure 1.**
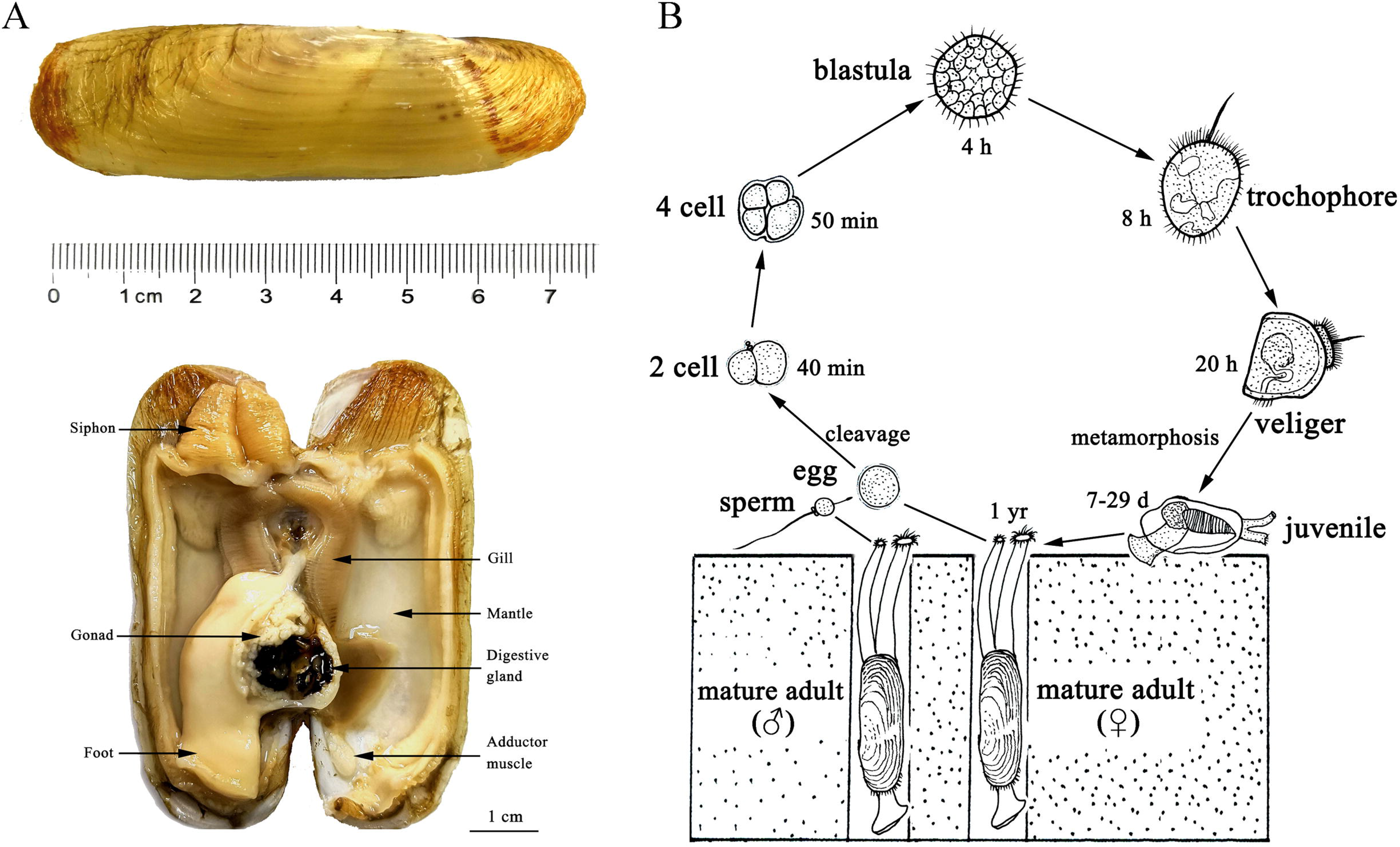
**A**. The appearance and anatomic structures of an adult razor clam. **B**. A pelago-benthic life cycle of the razor clam.

As living in estuarine and intertidal region, the razor clam faces tremendous exposure to extreme environmental stresses such as drastic salinity fluctuation, highly variable temperature, high concentration of ammonia nitrogen and hydrogen sulfide. Unlike oysters, mussels and most clams with thick and sealed shells for protecting their soft bodies, the razor clam with two thin and unclosed shells has chosen a survival strategy of deep-burrowing lifestyle with high tolerance of a broad range of salinities (5-45‰), making it an ideal model to investigate the adaptive mechanisms of deep-burrowing lifestyle. Despite its economic impact and representative of the less-understood deep-burrowing bivalve lifestyle, there are few genomic resources for exploring its unique biology and adaptive evolution. Here, we generated the high-quality chromosomal-level genome assembly and comprehensive transcriptomes of *S. constricta* and investigated the transcriptomic changes under different environmental stresses. These genomic resources will lay a prime foundation for future studies of its lifestyle-related adaptive evolution and genetic improvement in commercial breeding programs.

### Genomic DNA preparation, PacBio and Illumina sequencing

An individual *S. constricta* was collected from the brood stock at the genetic breeding research center of Zhejiang Wanli University. Genomic DNA was extracted from muscle tissues using a phenol-chloroform method as described in the protocol (Green and Sambrook, 2012). High molecular weight genomic DNA was sheared into fragments of ~30 kb using a Covaris ultrasonicators (Covaris, Woburn, MA, USA). The fragments were enzymatically repaired and converted into SMRTbell™ template library following the manufacturer’s instructions. Size-selection was performed to enrich the DNA fragments longer than 10 kb for sequencing on a Pacific Biosciences (PacBio) Sequel Single Molecule Real Time (SMRT) platform. The genomic library was sequenced in 6 cells, generating 10,549,576 subreads with a N50 length of 13,619 bp, and accounting for a total of 101.79 Gb. A paired-end Illumina library with an insert size of 300 bp were prepared with an Illumina Genomic DNA sample Preparation kit and sequenced on an Illumina Xten system, yielding a total of 129.73 Gb reads with an insert size of 350 bp (Supplementary Table S1).

### Estimation of the genome size and sequencing coverage

The Illumina short reads were first trimmed to remove adaptors and reads with more than 10% ambiguous or more than 20% low-quality bases using Trimmomatic (Bolger et al., 2014). The distribution of 17-mer frequency was estimated using the clean reads. A total of 10^10^ k-mer was identified with the peak depth of coverage being 80. Based on the formula: genome size = k-mer number/peak depth (Varshney et al., 2011), the genome size of *S. constricta* was estimated to be 1,244.27 Mb, with a heterozygous ratio of 1.53% and repeat rate of 53.12% (Supplementary Figure S1).

### *De novo* genome assembly and quality assessment

PacBio long reads were corrected and assembled using the Falcon package (Chin et al., 2016). Briefly, all the raw reads yielded by Pacbio platform were aligned to each other to identify overlaps with DALIGNER. The overlap data and raw subreads were then processed for consensus calling. After the error-correction, overlaps were detected in the preassembled error-corrected reads and used to construct a directed fragment assembly string graph. Contigs were constructed by finding the paths from the string graph. The consensus calling of preceding assembly was performed with Quiver. Subsequently, the paired-end clean reads yielded by Illumina platform were aligned to polish the assembly using Pilon (Walker et al., 2014). The resulting assembly consisted of 10,981 contigs, comprising 1,331.97 Mb with a contig N50 of 678,857 bp and GC contents of 35.46% (Table 1).

**Table 1.**
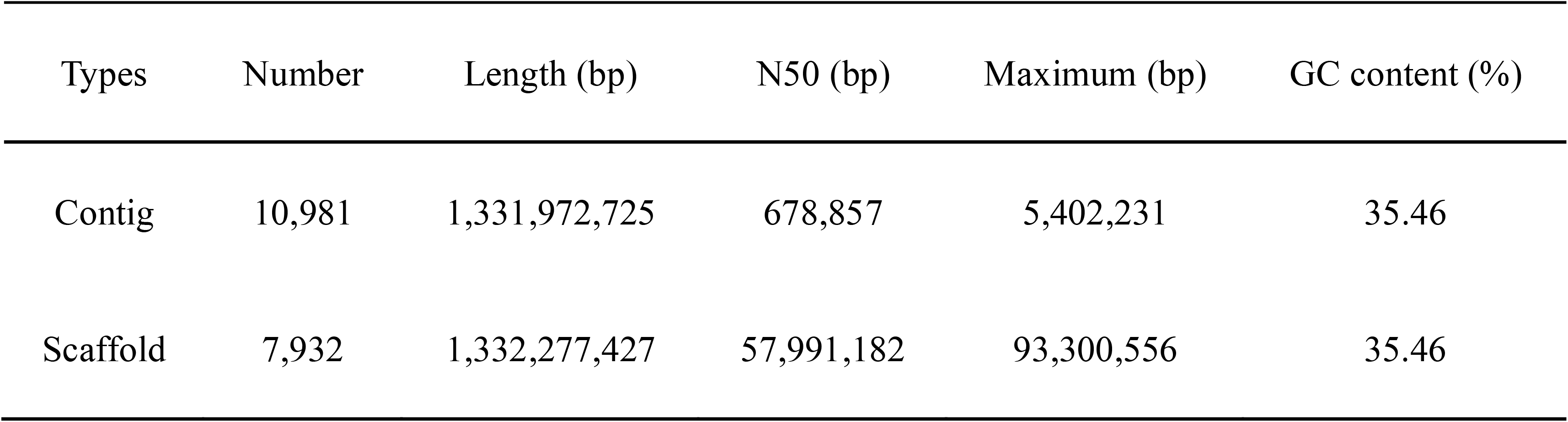
Statistics of the genome assembly of *Sinonovacula constricta*

| Types | Number | Length (bp) | N50 (bp) | Maximum (bp) | GC content (%) |
| --- | --- | --- | --- | --- | --- |
| Contig | 10,981 | 1,331,972,725 | 678,857 | 5,402,231 | 35.46 |
| Scaffold | 7,932 | 1,332,277,427 | 57,991,182 | 93,300,556 | 35.46 |

To assess the integrity of the genome assembly, Illumina short-insert library reads were mapped to the contigs using BWA (version 0.6.2). In summary, 88.90% of the assembled genome sequences were covered by 93.93% of the total reads (Supplementary Table S2). The genome completeness was also evaluated using both Core Eukaryotic Genes Mapping Approach (CEGMA) analysis (Parra et al., 2007) and Benchmarking Universal Single-Copy Orthologs (BUSCO version 3) analysis (Waterhouse et al., 2017). The CEGMA analysis identified 227 of the 248 core eukaryotic genes (91.53%), and the BUSCO analysis unveiled 868 of the 978 near-universal single-copy metazoan orthologs (88.7%), indicating a high integrity of the genome assembly (Supplementary Table S3 and S4).

### Illumina transcriptome sequencing and analysis

Transcriptomic samples from different developmental stages and different adult tissues were collected and sequenced for genome annotation. Embryos at four developmental stages (eggs, four cells, blastulae, gastrulae), and larvae at four developmental stages (trochophore larvae, D-shaped larvae, umbo larvae, and juvenile) were collected at the hatchery of genetic breeding research center of Zhejiang Wanli University. Artificial fertilization and larval culture were performed as previously described (Dong et al., 2012). For each developmental stage, over 1,000 individuals were collected for RNA extraction. Eight tissues (Figure 1A), including gill, digestive gland, foot, mantle, adductor muscle, siphon, gonad (testis and ovary) were dissected from one to three adult individuals and stored at −80°C after flash frozen in liquid nitrogen.

Transcriptomic samples under salinities of 3‰, 25‰, and 38‰ were collected and sequenced to identify genes and pathways involved in salt tolerance. The *S. constricta* were subjected to salt stress for 16 hours under 22°C at extreme concentration of low-salinity (3‰) and high-salinity (38‰) with the control concentration of normal-salinity (25‰). Three replicate tanks for each group were set and each replicate included 10 individuals. For the low-salinity group, the salinity was deceased 3‰ per hour though pouring into fresh water to target salinity of 3‰, and then maintained for 16 hours. For the high-salinity groups, the salinity was raised 2‰ per hour though pouring into artificial sea water to target salinity of 38‰, and then maintained for 16 hours. Gills were dissected from three individuals of each replicate and stored at −80°C.

Total mRNA was extracted from all the collected samples with TRIzol reagent (OMEGA, America) following the manufacturer’s instructions. A paired-end Illumina library was constructed for each sample with an insert size of 300 bp and sequenced on an Illumina X Ten system. Around 5-7 Gb of paired-end raw reads were yielded for each sample. Clean reads were obtained by removing reads containing adapter, reads containing ploy-N and low-quality reads by Trimmomatic (Bolger et al., 2014) (Supplementary Table S5). The clean reads were aligned to the indexed *S. constricta* reference genome using Hisat2 version 2.0.5 (Kim et al., 2015). The clean reads in the samples of different adult tissues and salt stress were mapped onto the reference genome with high proportion of around 70-80%, while samples of different development stages with relative low proportion because of mixed thousand individuals increasing the high SNP heterozygosity (Supplementary Table S6). The featureCounts version 1.5.0 (Liao et al., 2014) was used to count the reads numbers mapped to each gene and the gene expression level was calculated as FPKM (Fragments Per Kilobase of transcript sequence per Millions base pairs) for each gene.

The identification of differentially expressed genes (DEGs) between different salinity groups was performed using the DESeq2 R package version 1.16.1 with an adjusted P-value <0.05 (Love et al., 2014). The numbers of up- and down-regulated DEGs were 462 and 655 between the high-salinity group versus the normal-salinity group, respectively while the numbers of up- and down-regulated DEGs were 898 and 826 between the low-salinity group versus the normal-salinity group, respectively (Supplementary Figure S2). Gene Ontology (GO) enrichment analysis of DEGs was implemented by the clusterProfiler R package with corrected P-value <0.05 considered significantly enriched GO terms (Yu et al., 2012). The clusterProfiler R package is also used to test the statistical enrichment of DEGs in KEGG pathways (Yu et al., 2012). The GO enrichment results demonstrated that the DEGs were significantly enriched in the biological processes of transmembrane transport (GO:0055085) and aminoglycan metabolic process (GO:0006022) and the molecular functions of transmembrane transporter activity (GO:0022857) and (Supplementary Figure S3). The KEGG pathway enrichment results indicated that the DEGs were significantly enriched in amino acid metabolic pathways such as glycine, serine and threonine metabolism, and arginine and proline metabolism, and the energy metabolic pathways such as glycolysis/gluconeogenesis and citrate cycle (Supplementary Figure S4).

### Full-length transcriptome sequencing and analysis

Full-length RNA sequencing was also performed using the mixed RNAs from the samples of eight development stages and eight adult tissues. Three libraries with different insert lengths, e.g. 1-2k, 2-3k, and 3-6k, were constructed and sequenced on a PacBio Sequel platform. A total of 1,064,194 post-filter polymerase reads were obtained from 7 SMRT cells, including 688,944 full-length non-chimeric reads (Supplementary Table S7). The full-length RNA transcriptomic analysis was performed with the SMRT Link v4.0.0 software suite (https://www.pacb.com/support/software-downloads). After redundant sequences clustering using the ICE (Iterative Clustering and Error correction) algorithm, consensus sequences building using the pbdagcon tool with DAGCon (Directed Acyclic Graph Consensus) algorithm, and consensus sequence polishing with Quiver, a total of 61,620 high-quality (>0.99) and 358,297 low-quality (<0.99) transcript sequences were obtained. Then, the transcript sequences were polished and corrected using Illumina reads with LoRDEC (Salmela and Rivals, 2014). Finally, the corrected transcripts were further clustered with CD-HIT (version 4.6) (Li and Godzik, 2006), resulting in 75,225 Unigenes and 276,484 transcript isoforms. The full-length transcripts were further used to annotate the protein-coding genes in the genome as the direct evidences. The statistical information for full-length transcriptome analysis is listed in Supplementary Table S7.

### Repetitive sequence annotation

Repetitive sequences in the genome assembly were identified through *ab initio* prediction and homology-based searches. RepeatScout (version 1.0.5) and Repeat Modeler version 1.0.11 (http://www.repeatmasker.org) were used for *de novo* identification of repeat families in the *S. constricta* genome. Full length long terminal repeat (LTR) retrotransposons were also identified using the LTR-finder (version 1.0.2) (Xu and Wang, 2007) with the parameters “-E -C”. Tandem Repeats Finder (TRF version 4.09) (Benson, 1999) was used to screen tandem repeats with the parameters “match=2, mismatching penalty=7, indel penalty=7, match probability=80, indel probability=10, minimum alignment score=50, maximum period size=500”. The predicted repetitive sequences along with the RepBase database (Bao et al., 2015) were used for homology-based searches using Repeatmasker (version 4.5.0) with the parameters “-a -nolow -no_is -norna -parallel 32 -small -xsmall -poly -e ncbi -pvalue 0.0001” (Tarailo-Graovac and Chen, 2009).

Finally, a total of 675,404,889 bp repetitive sequences were identified, accounting for 50.71% of the assembled genome (Table 2), which is consistent with our genome survey result of 53.12%. Repetitive sequences were dominated by tandem repeats (15.39%) and followed by DNA transposons (14.38%) and LTR retrotransposons (10.84%) (Table 2).

**Table 2.** Statistics of the repetitive sequences

| Type | Repeat size (bp) | % of genome |
| --- | --- | --- |
| DNA | 191,499,094 | 14.38 |
| LINE | 71,938,692 | 5.4 |
| LTR | 144,451,530 | 10.84 |
| SINE | 5,528,172 | 0.42 |
| Tandem repeat | 204,889,587 | 15.38 |
| Other | 157,232 | 0.01 |
| Unknown | 56,940,582 | 4.27 |
| Total | 675,404,889 | 50.71 |

### Protein-coding gene prediction and annotation

Gene annotation was performed based on *de novo* prediction, homology-based searches, and transcriptome assisted methods. Protein sequences of Yesso scallop (*Patinopecte yessoensis*), Pacific oyster (*Crassostrea gigas*), owl limpet (*Lottia gigantea*), octopus (*Octopus bimaculoides*), leech (*Helobdella robusta*), nematode (*Caenorhabditis elegans*), fruit fly (*Drosophila melanogaster*), sea urchin (*Strongylocentrotus purpuratus*), ascidian (*Ciona intestinalis*), Florida lancelet (*Branchiostoma floridae*), and human (*Homo sapiens*) were downloaded from NCBI and aligned to the genome assembly using TBLASTN with the parameters “-evalue 1e-5”. The gene structures were predicted with GeneWise (version 2.4.1) (Doerks et al., 2002). The Illumina RNA-seq reads of the eight tissues and eight developmental stages were aligned to the genome assembly using Tophat (version 2.1.1) (Trapnell et al., 2009). Cufflinks (version 2.1.0) (Trapnell et al., 2010) was used to generate gene models with the parameter “-multi-read-correct”. The resulting GTF file along with the PacBio Iso-seq transcripts was utilized to model gene structures with the PASA pipeline (version 2.0.2) (Haas et al., 2008).

Five *de novo* gene prediction packages, including Augustus (version 2.5.5) (Stanke et al., 2006), glimmerHMM (version 3.01) (Majoros et al., 2004), SNAP (version 2006-07-28) (Leskovec and Sosic, 2016), Geneid (version 1.4) (Parra et al., 2000), and Genscan (version 3.1) (Burge and Karlin, 1997) were used to predict genes with the repeat-masked genome sequences by default settings. All the gene model evidences were integrated using EVidenceModeler (version 1.1.1) (Haas et al., 2008). Finally, 26,273 protein-coding genes were identified in the *S. constricta* genome (Supplementary Table S8).

The functional annotations were performed by aligning the predicted protein sequences to public databases including KEGG, SwissProt and NCBI-NR databases using BLASTP with the E-value threshold of 1e-5. InterProScan (v.4.8) (Jones et al., 2014) was also used to identify motifs and domains by searching the Pfam, InterPro and Gene Ontology (GO) databases. Taken together, 26,140 (99.5%) of the 26,273 genes could be annotated by at least one database (Table 3).

**Table 3.** Statistics of gene annotation to different databases

| Annotation database | Number of annotated genes | Percentage (%) |
| --- | --- | --- |
| NR | 23,844 | 90.88 |
| Swiss-Prot | 18,131 | 69.1 |
| KEGG | 18,928 | 72.14 |
| InterProScan | 25,475 | 97.1 |
| Pfam | 15,391 | 58.66 |
| GO | 22,956 | 87.49 |
| Total | 26,140 | 99.50 |

### Noncoding RNA prediction and annotation

The noncoding RNA genes, including miRNAs, rRNAs, snRNAs, and tRNAs, were annotated in the *S. constricta* genome. tRNAs were predicated by tRNAscan-SE 2.0 (Lowe and Chan, 2016) with eukaryote parameters. The miRNAs and snRNAs were screened using INFERNAL 1.1.2 against the Rfam database (version 14.1) (Kalvari et al., 2018) with default parameters. Finally, 968 miRNAs, 3,354 tRNAs, 822 rRNAs, and 298 snRNAs were identified (Supplementary Table S9).

### Gene family and phylogenetic analysis

Fifteen Eumetazoa species were selected for gene family analysis, including *H. sapiens*, *B. floridae*, *D. melanogaster*, European honey bee (*Apis mellifera*), Californian leech (*Helobdella robusta*), ocean-dwelling worm (*Capitella teleta*), *O. bimaculoides*, *L. gigantea*, California sea hare (*Aplysia californica*), *C. gigas*, American oyster (*Crassostrea virginica*), Sydney rock oyster (*Saccostrea glomerata*), *P. yessoensis*, Chinese scallop (*Chlamys farreri*), and Starlet sea anemone (*Nematostella vectensis*). All data were downloaded from either NCBI or Ensembl. The longest protein sequence was selected to represent the gene with multiple alternative splicing isoforms. Gene family clusters from all the 16 species were first assigned using OrthoMCL (version 2.0.9) (Li et al., 2003) with an inflation value of 1.5. CAFE (version 3) (De Bie et al., 2006) was used to analyze gene family expansion and contraction under maximum likelihood framework. The protein-coding genes from all the 16 species were assigned into 39,058 families with 337 strict single-copy orthologs. In the *S. constricta* genome, a total of 12,945 gene families were identified, 803 of which were specifically possessed by *S. constricta.* Compared with the other 15 species, *S. constricta* has 193 expanded and 31 contracted gene families (Figure 2). Notably, cytoskeletal protein alpha tubulin (*TUA*) family and motor protein dynein heavy chain (*DYH*) family are rapidly expanded in the *S. constricta* genome (Figure 3A). They play vital roles in the microtubule architecture and the bending movement of cilia (Mohri et al., 2012). The razor clam has a well-developed ciliary system for gill filtering, food-particles retaining, and water pumping (Morton, 1984). The adjoining cilia generate effective beat through coordinated wavelike movements. The pumping rate of the ciliary system in the gill and mantle cavity can be adjusted to generate powerful currents to facilitate the principal sorting and retaining of suspended particles in the labial palps. Effluxes can also be ejected from the pedal gape to flush away sources of irritation detected by the sensory tentacles (Morton, 1984). The transcriptomic data revealed that the *TUA* and *DYH* genes are highly expressed in the gills (Figure 3B and 3C), suggesting that the expansion of these genes could be an adaptation to the deep-burrowing lifestyle.

**Figure 2.**
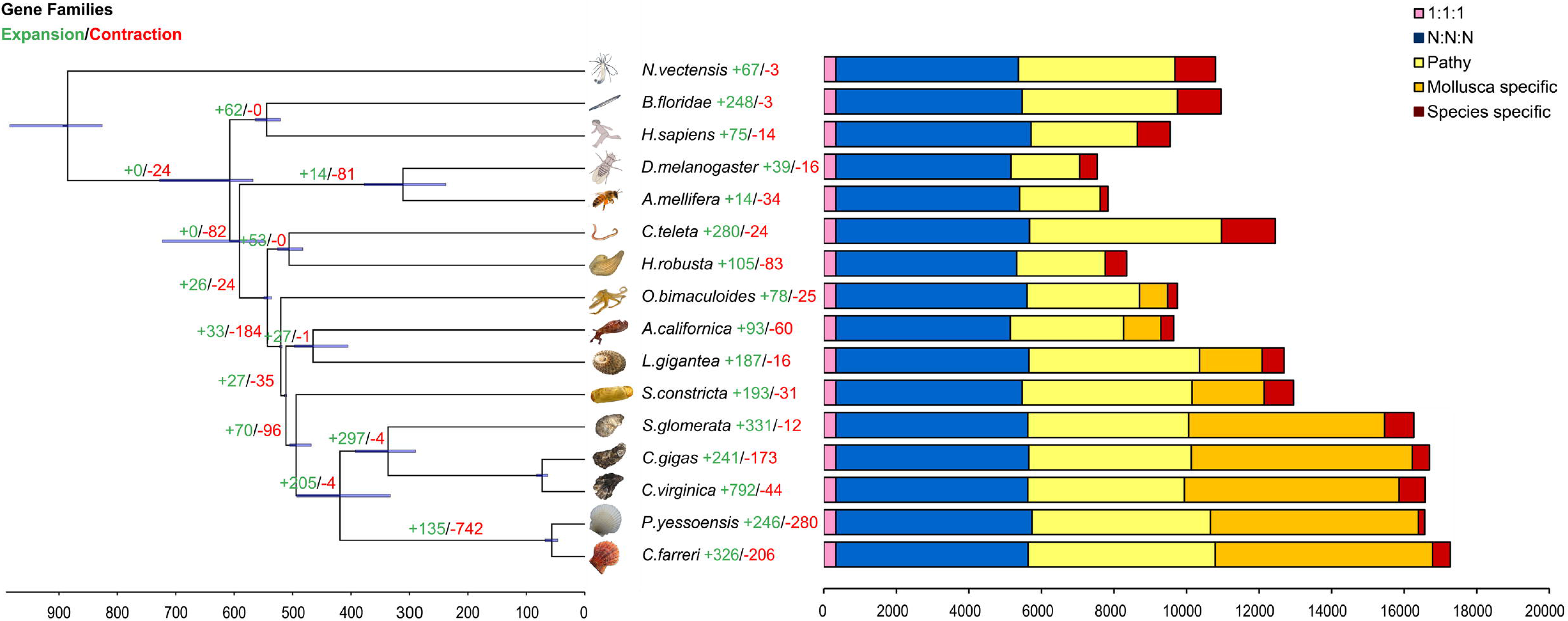
Phylogenetic tree and number of shared orthologs among *S. constricta* and other animal species. Numbers of gene families undergoing expansion and contraction for each lineage are exhibited as red and green, respectively.

**Figure 3.**
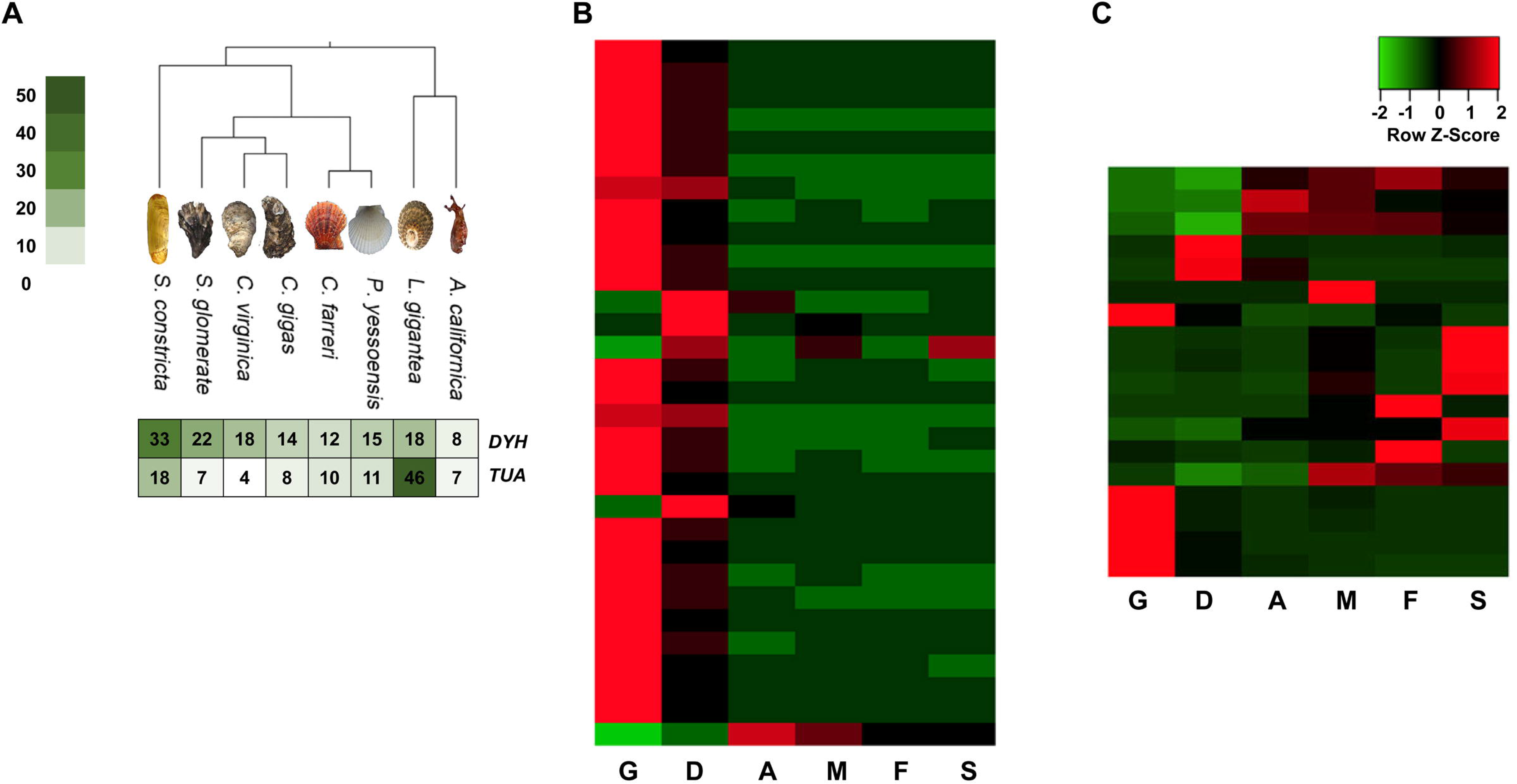
**A.**The comparison of the copy numbers of dynein heavy chain (*DYH*) and alpha tubulin (TUA) genes in 8 molluscan species. **B & C.** The tissue-wide expression patterns of *DYH* genes and *TUA* genes. Abbreviations: G, gill; D, digestive gland; A. adductor muscle; M, mantle; F, foot; S, siphon.

Phylogenetic inference of the 16 species was performed with the 337 single-copy orthologs. Multiple sequences alignment was conducted for the protein sequences of each ortholog gene using MUSCLE (version 3.8.31) (Edgar, 2004) separately. The alignments for all the orthologs were then concatenated into a super alignment matrix with 241,349 amino acids. RAxML (version 8.2.12) (Stamatakis, 2014) was used to infer the alignment matrix by a maximum likelihood method with the substitution model PROTGAMMAAUTO. Bootstrapping with 100 replicates was used for node support. Divergence time between species was estimated using MCMCTree in PAML package (version 4.7a) (Yang, 1997) with the parameters of “burn-in = 1,000, sample-number = 1,000,000, sample-frequency = 2”. The constructed maximum likelihood phylogenetic tree revealed that *S. constricta* clustered with other bivalve species and diverged ~494 million years ago (Mya) from the lineage leading to oysters and scallops (Figure 2).

### Hi-C scaffolding and macro-synteny analysis

Adductor muscle tissue of a razor clam from the same population was collected for Hi-C library construction. The tissue specimen was fixed with 1% formaldehyde and the genomic DNA was cross-linked, digested by restriction enzymes HindIII, labeled with biotinylated residue, and end repaired. The library was sequenced on an Illumina NovaSeq platform, generating 156.73 G of raw reads. The raw reads were truncated at the junctions and aligned to the polished genome using BWA (version 0.7.17) with default parameters. Only the unique aligned reads with a mapping quality over 20 were further processed. After filtering invalid interaction pairs by HiC-Pro (v.2.8.0) (Servant et al., 2015), 30.32% of the clean reads were valid pairs and utilized to evaluate the interaction strength among whole genome contigs. Lachesis (version 2e27abb) was used to cluster and anchor the contigs to the chromosomes using an agglomerative hierarchical clustering method (Burton et al., 2013). Finally, 3,068 contigs, accounting for 87.82% of the total bases, were clustered into 19 linkage groups (Figure 4A), which was consistent with the karyotype revealed by previous studies (Wang et al., 1998). The ancient ortholog genes exhibited remarkable preservation of ancestral bilaterian linkage groups (Simakov et al., 2013; Wang et al., 2017) with a conservation index (CI) of 0.71, indicating the considerable accuracy of the Hi-C clustering (Figure 4B).

**Figure 4.**
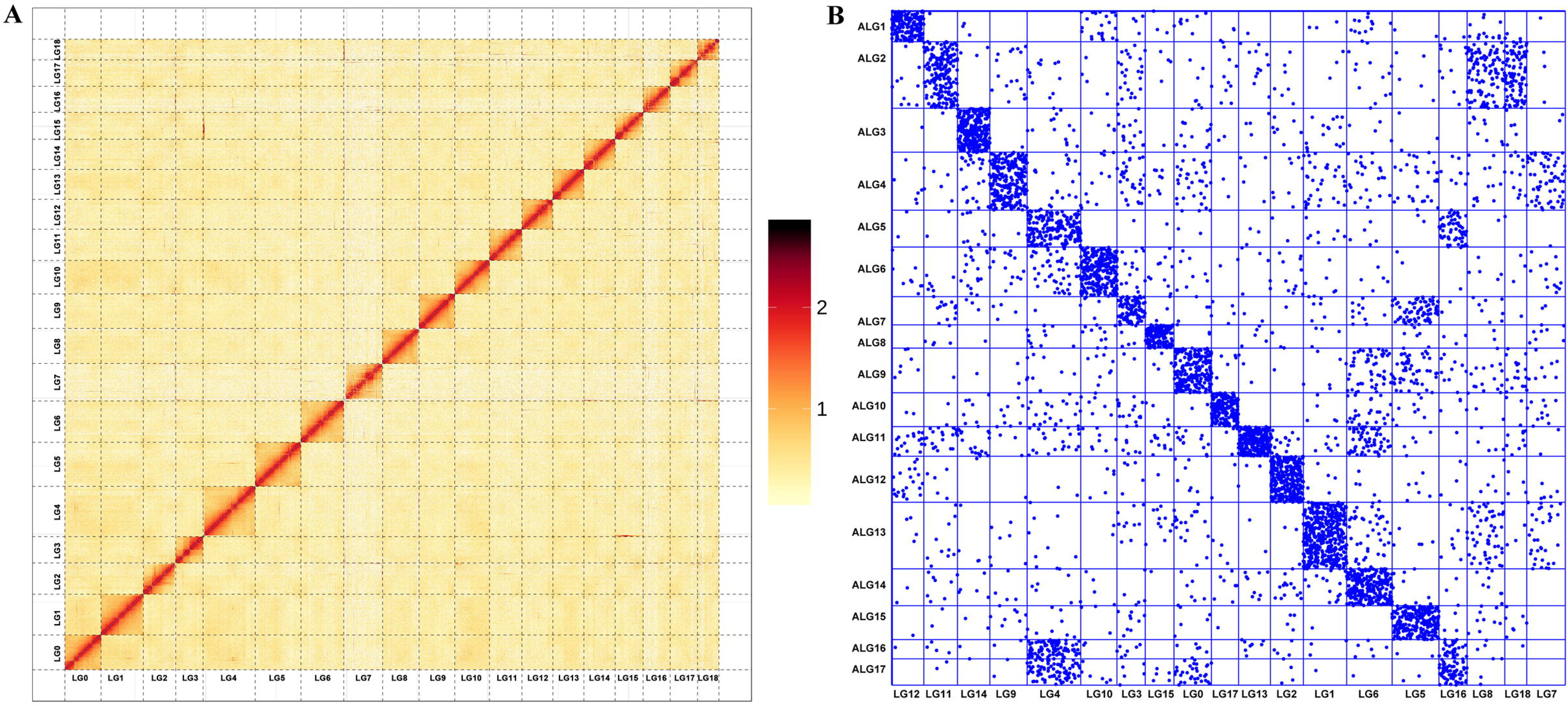
**A.** Hi-C interaction heat map of of *S. constricta*. **B.** Chromosome-based macro-synteny between *S. constricta* and the 17 presumed bilaterian ALGs retrieved from Simakov et al. (2013).

## Conclusions

We assembled a high-quality chromosomal-level reference genome of *S. constricta*, the first genome of the family Solenidae, along with comprehensive transcriptomic data of whole-ontogeny developmental stages and all major tissues (under normal and stressed conditions). The genomic and transcriptomic resources reported here would lay a prime foundation for future studies to elucidate the razor clam’ adaptive traits relating to deep-burrowing lifestyle (e.g., thin shells, advanced ciliary and siphon system, , extraordinary adaption to broad-range salinity and high concentration of ammonia nitrogen and hydrogen sulfide) and genetic improvement in commercial breeding programs.

## Ethics approval and consent to participate

All experimental procedures were approved by the Institutional Animal Care and Use Committee (IACUC) of Zhejiang Wanli University, China. All participants consent to publish the work under the “Consent to publish” heading.

## Data availability

The *S. constricta* genome assembly is available at NCBI (BioProject: PRJNA559038). RNA sequencing data files are available through the NCBI Sequence Read Archive (BioProject: PRJNA559056). The *S. constricta* genome assembly and annotation files also could be downloaded from the website http://202.121.66.128/clam-genome/zwu.htm.

## Acknowledgements

This work was financially supported by the National Key Research and Development Program of China (No.2018YFD0901405), Zhejiang Major Program of Science and Technology (No.2016C02055-9), Modern Agro-industry Technology Research System (No. CARS-49).

## Author contributions

Y. D., Z. L. and Z. B. conceived the project. H. Y., W. R. and L.H. conducted the environmental stress and collected the samples. Q. Z., S. W., J. R. and L. L. performed the genome assembly, annotation, transcriptome analysis and other bioinformatics analysis. Y. D., J. R., Q. Z., S. W. and Q. X. wrote and revised the manuscript. All authors read and approved the final manuscript.

## Supplementary materials

**Table S1.** Summary of the genomic sequencing reads

**Table S2.** Statistics of Illumina short reads coverage

**Table S3.** Summary genomic completeness by CEGMA

**Table S4.** Summary genomic completeness by BUSCO

**Table S5.** Summary of Illumina transcriptome sequencing data

**Table S6.** Summary of clean reads mapping

**Table S7.** Summary of PacBio full-length transcriptome sequencing

**Table S8.** Summary of the gene prediction results

**Table S9.** Summary of the non-coding RNA annotation

**Figure S1.** Genome survey of *Sinonovacula constricta* using 17-mer analysis

**Figure S2.** Volcano map of differentially expressed genes

**Figure S3.** Dot plot of GO enrichment of differentially expressed genes

**Figure S4.** Dot plot of KEGG pathway enrichment of differentially expressed gen

## Reference

Bao, W., Kojima, K.K., and Kohany, O.(2015). Repbase Update, a database of repetitive elements in eukaryotic genomes. Mobile DNA 6, 11.

Benson, G.(1999). Tandem repeats finder: a program to analyze DNA sequences. Nucleic acids research 27, 573–580.

Bolger, A.M., Lohse, M., and Usadel, B.(2014). Trimmomatic: a flexible trimmer for Illumina sequence data. Bioinformatics 30, 2114–2120.

Burge, C., and Karlin, S.(1997). Prediction of complete gene structures in human genomic DNA. Journal of molecular biology 268, 78–94.

Burton, J.N., Adey, A., Patwardhan, R.P., Qiu, R., Kitzman, J.O., and Shendure, J.(2013). Chromosome-scale scaffolding of de novo genome assemblies based on chromatin interactions. Nature biotechnology 31, 1119–1125.

Chin, C.S., Peluso, P., Sedlazeck, F.J., Nattestad, M., Concepcion, G.T., Clum, A., Dunn, C., O’Malley, R., Figueroa-Balderas, R., Morales-Cruz, A., et al.(2016). Phased diploid genome assembly with single-molecule real-time sequencing. Nature methods 13, 1050–1054.

De Bie, T., Cristianini, N., Demuth, J.P., and Hahn, M.W.(2006). CAFE: a computational tool for the study of gene family evolution. Bioinformatics 22, 1269–1271.

Doerks, T., Copley, R.R., Schultz, J., Ponting, C.P., and Bork, P.(2002). Systematic identification of novel protein domain families associated with nuclear functions. Genome research 12, 47–56.

Dong, Y.H., Yao, H.H., Zhang, P.Y., Shen, P.Y., Liiu, H.M., and Lin, Z.H.(2012). Cytological observation on fertilization and early cleavage in Sinonovaula constricta. Journal of fisheries of China 36, 1400–1409.

Edgar, R.C.(2004). MUSCLE: multiple sequence alignment with high accuracy and high throughput. Nucleic acids research 32, 1792–1797.

FAO(2018). FAO yearbook. Fishery and Aquaculture Statistics 2016. http://www.faoorg//fishery/publications/yearbooks/en.

Green, M., and Sambrook, J.(2012). Molecular Cloning: A Laboratory Manual. 4th Edition, Vol. II, Cold Spring Harbor Laboratory Press, New York.

Haas, B.J., Salzberg, S.L., Zhu, W., Pertea, M., Allen, J.E., Orvis, J., White, O., Buell, C.R., and Wortman, J.R.(2008). Automated eukaryotic gene structure annotation using EVidenceModeler and the Program to Assemble Spliced Alignments. Genome biology 9, R7.

Jones, P., Binns, D., Chang, H.Y., Fraser, M., Li, W., McAnulla, C., McWilliam, H., Maslen, J., Mitchell, A., Nuka, G., et al.(2014). InterProScan 5: genome-scale protein function classification. Bioinformatics 30, 1236–1240.

Kalvari, I., Nawrocki, E.P., Argasinska, J., Quinones-Olvera, N., Finn, R.D., Bateman, A., and Petrov, A.I.(2018). Non-Coding RNA Analysis Using the Rfam Database. Current protocols in bioinformatics 62, e51.

Kim, D., Langmead, B., and Salzberg, S.L.(2015). HISAT: a fast spliced aligner with low memory requirements. Nature methods 12, 357–360.

Leskovec, J., and Sosic, R.(2016). SNAP: A General Purpose Network Analysis and Graph Mining Library. ACM transactions on intelligent systems and technology 8.

Li, L., Stoeckert, C.J., Jr., and Roos, D.S.(2003). OrthoMCL: identification of ortholog groups for eukaryotic genomes. Genome research 13, 2178–2189.

Li, W., and Godzik, A.(2006). Cd-hit: a fast program for clustering and comparing large sets of protein or nucleotide sequences. Bioinformatics 22, 1658–1659.

Liao, Y., Smyth, G.K., and Shi, W.(2014). featureCounts: an efficient general purpose program for assigning sequence reads to genomic features. Bioinformatics 30, 923–930.

Love, M.I., Huber, W., and Anders, S.(2014). Moderated estimation of fold change and dispersion for RNA-seq data with DESeq2. Genome biology 15, 550.

Lowe, T.M., and Chan, P.P.(2016). tRNAscan-SE On-line: integrating search and context for analysis of transfer RNA genes. Nucleic acids research 44, W54–57.

Majoros, W.H., Pertea, M., and Salzberg, S.L.(2004). TigrScan and GlimmerHMM: two open source ab initio eukaryotic gene-finders. Bioinformatics 20, 2878–2879.

Mohri, H., Inaba, K., Ishijima, S., and Baba, S.A.(2012). Tubulin-dynein system in flagellar and ciliary movement. Proceedings of the Japan Academy Series B, Physical and biological sciences 88, 397–415.

Morton, B.(1984). The functional morphology of Sinonovucufu constvictu with a discussion on the taxonomic status of the Novaculininae (Bivalvia). J Zool, Lond 202, 299–325.

Parra, G., Blanco, E., and Guigo, R.(2000). GeneID in Drosophila. Genome research 10, 511–515.

Parra, G., Bradnam, K., and Korf, I.(2007). CEGMA: a pipeline to accurately annotate core genes in eukaryotic genomes. Bioinformatics 23, 1061–1067.

Salmela, L., and Rivals, E.(2014). LoRDEC: accurate and efficient long read error correction. Bioinformatics 30, 3506–3514.

Servant, N., Varoquaux, N., Lajoie, B.R., Viara, E., Chen, C.J., Vert, J.P., Heard, E., Dekker, J., and Barillot, E.(2015). HiC-Pro: an optimized and flexible pipeline for Hi-C data processing. Genome biology 16, 259.

Simakov, O., Marletaz, F., Cho, S.J., Edsinger-Gonzales, E., Havlak, P., Hellsten, U., Kuo, D.H., Larsson, T., Lv, J., Arendt, D., et al.(2013). Insights into bilaterian evolution from three spiralian genomes. Nature 493, 526–531.

Stamatakis, A.(2014). RAxML version 8: a tool for phylogenetic analysis and post-analysis of large phylogenies. Bioinformatics 30, 1312–1313.

Stanke, M., Keller, O., Gunduz, I., Hayes, A., Waack, S., and Morgenstern, B.(2006). AUGUSTUS: ab initio prediction of alternative transcripts. Nucleic acids research 34, W435–439.

Trapnell, C., Pachter, L., and Salzberg, S.L.(2009). TopHat: discovering splice junctions with RNA-Seq. Bioinformatics 25, 1105–1111.

Trapnell, C., Williams, B.A., Pertea, G., Mortazavi, A., Kwan, G., van Baren, M.J., Salzberg, S.L., Wold, B.J., and Pachter, L.(2010). Transcript assembly and quantification by RNA-Seq reveals unannotated transcripts and isoform switching during cell differentiation. Nature biotechnology 28, 511–515.

Varshney, R.K., Chen, W., Li, Y., Bharti, A.K., Saxena, R.K., Schlueter, J.A., Donoghue, M.T., Azam, S., Fan, G., Whaley, A.M., et al.(2011). Draft genome sequence of pigeonpea (Cajanus cajan), an orphan legume crop of resource-poor farmers. Nature biotechnology 30, 83–89.

Walker, B.J., Abeel, T., Shea, T., Priest, M., Abouelliel, A., Sakthikumar, S., Cuomo, C.A., Zeng, Q., Wortman, J., Young, S.K., et al.(2014). Pilon: an integrated tool for comprehensive microbial variant detection and genome assembly improvement. PloS one 9, e112963.

Wang, J.X., Zhao, X.F., Zhou, L.H., and Xiang, J.H.(1998). Chromosome study of Sinonovacula constricta (Bivalvia) Oceanologia Et Limnologia Sinica 29, 191–196.

Wang, S., Zhang, J., Jiao, W., Li, J., Xun, X., Sun, Y., Guo, X., Huan, P., Dong, B., Zhang, L., et al.(2017). Scallop genome provides insights into evolution of bilaterian karyotype and development. Nature ecology & evolution 1, 120.

Waterhouse, R.M., Seppey, M., Simao, F.A., Manni, M., Ioannidis, P., Klioutchnikov, G., Kriventseva, E.V., and Zdobnov, E.M.(2017). BUSCO applications from quality assessments to gene prediction and phylogenomics. Molecular biology and evolution.

Xu, Z., and Wang, H.(2007). LTR_FINDER: an efficient tool for the prediction of full-length LTR retrotransposons. Nucleic acids research 35, W265–268.

Yang, Z.(1997). PAML: a program package for phylogenetic analysis by maximum likelihood. Computer applications in the biosciences : CABIOS 13, 555–556.

Yu, G., Wang, L.G., Han, Y., and He, Q.Y.(2012). clusterProfiler: an R package for comparing biological themes among gene clusters. Omics : a journal of integrative biology 16, 284–287.

