## Supplemental Figures for "The chromosomal-level genome assembly and comprehensive transcriptomes of Chinese razor clam (*Sinonovacula constricta*) with deep-burrowing life style and broad-range salinity adaptation"

**Supplementary material**

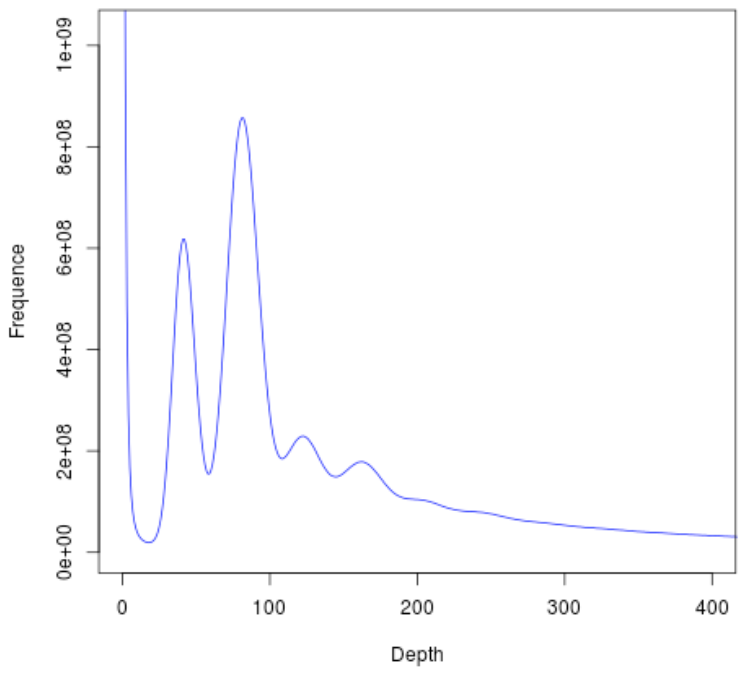

**Figure S1. Genome survey of *Sinonovacula constricta* using 17-mer analysis.**

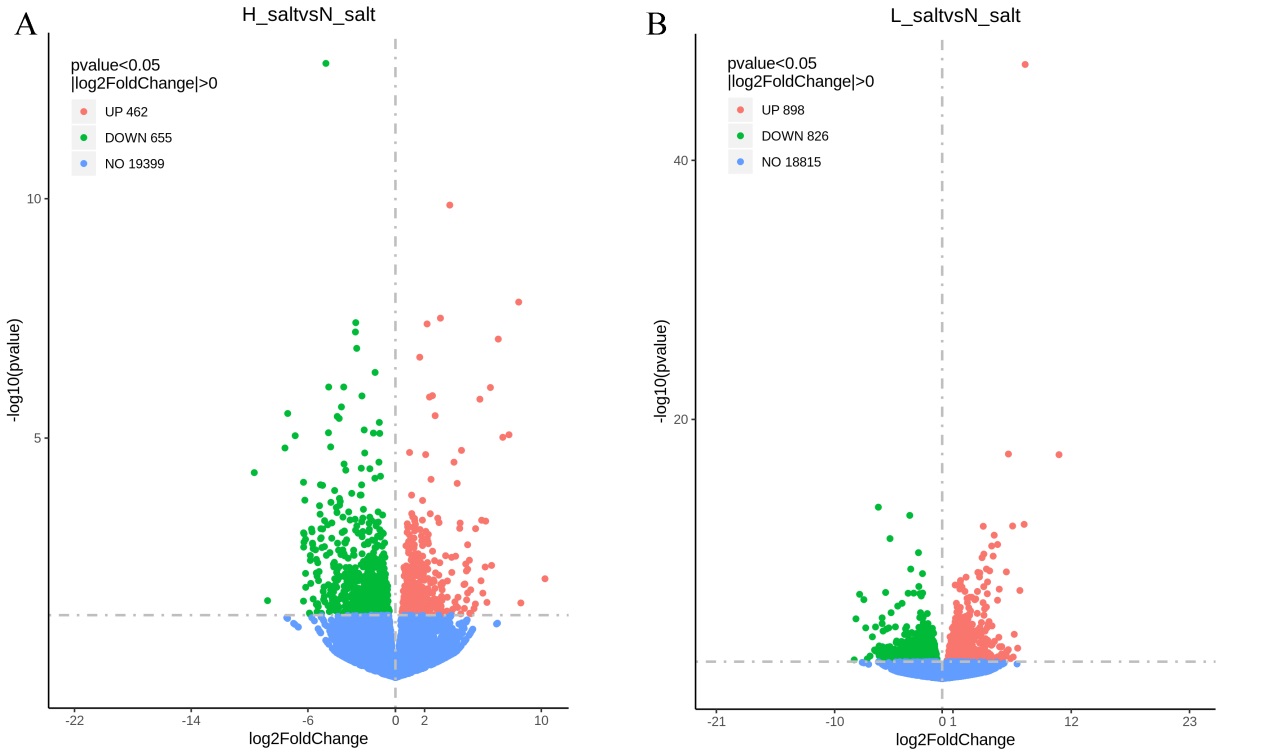

**Figure S2. Volcano map of differentially expressed genes**

A. Differential expressed genes between high salinity group and normal salinity group. B. Differential expressed genes between low salinity group and normal salinity group.

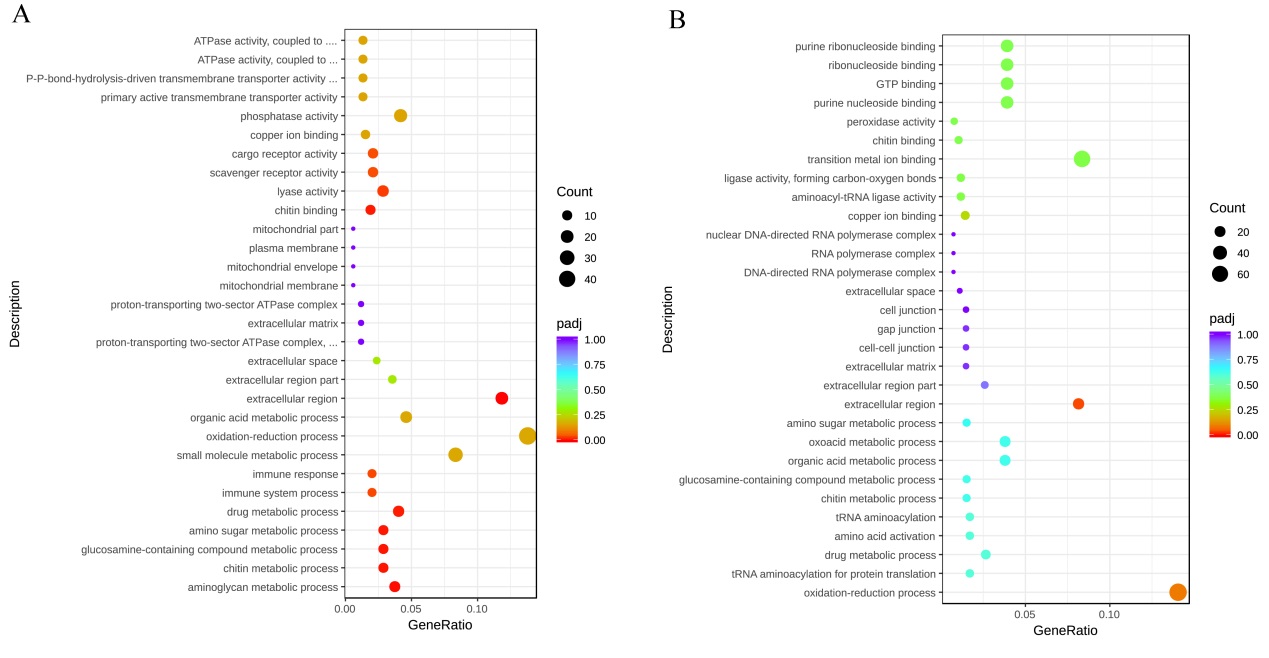

**Figure S3. Dot plot of GO enrichment of differentially expressed genes**

A. GO enrichment of differential expressed genes between high salinity group and normal salinity group. B. GO enrichment of differential expressed genes between low salinity group and normal salinity group.

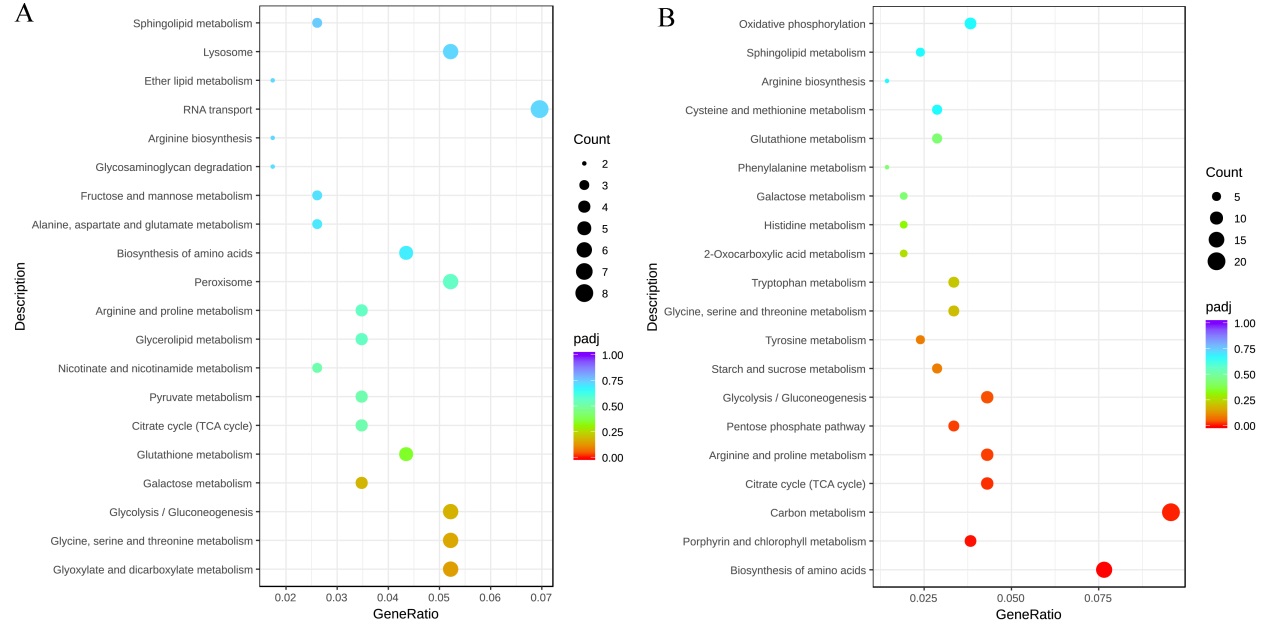
**Figure S4. Dot plot of KEGG pathway enrichment of differentially expressed genes**

A. KEGG pathway enrichment of differential expressed genes between high salinity group and normal salinity group. B. KEGG pathway enrichment of differential expressed genes between low salinity group and normal salinity group.
