## Supplemental Table for "The chromosomal-level genome assembly and comprehensive transcriptomes of Chinese razor clam (*Sinonovacula constricta*) with deep-burrowing life style and broad-range salinity adaptation"

**Supplementary material**

**Table S1. Summary of the genomic sequencing reads**

| Library | Total data (Gb) | Depth (×) |
| --- | --- | --- |
| Illumina reads | 129.73 | 104.62 |
| PacBio reads | 101.79 | 82.01 |
| Total | 231.52 | 186.63 |

**Table S2. Statistics of Illumina short reads coverage**

| Genome assembly | Parameter |
| --- | --- |
| Average sequencing depth | 87.11 |
| Mapping rate (%) | 93.93 |
| Coverage (%) | 88.90 |
| Coverage at least 4X (%) | 86.67 |
| Coverage at least 10X (%) | 85.18 |
| Coverage at least 20X (%) | 83.65 |

**Table S3. Summary genomic completeness by CEGMA**

| Number of core eukaryotic genes | Complete | | | Complete and Partial | | |
| --- | --- | --- | --- | --- | --- | --- |
|  | Number | | Percentage | Number | | Percentage |
| 248 | 198 | 79.84 | | 227 | 91.53 | |

**Table S4. Summary genomic completeness by BUSCO**

| Number of core metazoan genes | Complete | | Fragmented | Missing |
| --- | --- | --- | --- | --- |
|  | single copy | duplicated |  |  |
| 978 | 832 | 36 | 40 | 70 |

**Table S5. Summary of Illumina transcriptome sequencing data**

| Sample type | Sample name | Raw reads | Clean reads | Clean data (Gb) | Base quality  (Q20) |
| --- | --- | --- | --- | --- | --- |
| Different development stages | Egg | 43,935,046 | 42,881,284 | 6.43 | 97.28 |
|  | Four cells | 43,085,794 | 42,600,768 | 6.39 | 96.44 |
|  | Blastulae | 47,526,530 | 46,718,170 | 7.01 | 96.18 |
|  | Gastrulae | 43,935,046 | 42,881,284 | 6.43 | 97.28 |
|  | Trochophore | 43,935,046 | 42,881,284 | 6.43 | 97.28 |
|  | D-stage larvae | 36,806,136 | 36,163,274 | 5.42 | 95.99 |
|  | Umbo larvae | 43,492,564 | 43,044,208 | 6.46 | 96.62 |
|  | Juvenile | 44,296,144 | 43,739,942 | 6.56 | 96.30 |
| Different adult tissues | Gill | 58,314,622 | 52,571,902 | 5.31 | 97.71 |
|  | Foot | 57,279,662 | 51,774,716 | 5.23 | 97.75 |
|  | Adductor muscle | 50,833,380 | 45,661,534 | 4.61 | 97.66 |
|  | Digestive gland | 50,796,292 | 45,624,660 | 4.61 | 97.62 |
|  | Mantle | 54,234,040 | 48,455,080 | 4.89 | 97.51 |
|  | Siphon | 54,593,774 | 49,368,016 | 4.99 | 97.72 |
|  | Ovary01 | 34,903,986 | 34,468,792 | 4.33 | 98.05 |
|  | Ovary02 | 36,188,356 | 35,741,542 | 4.50 | 97.91 |
|  | Ovary03 | 36,718,496 | 36,258,132 | 4.56 | 97.95 |
|  | Testis01 | 32,433,246 | 32,024,558 | 4.03 | 97.85 |
|  | Testis02 | 28,685,592 | 28,336,062 | 3.56 | 97.99 |
|  | Testis03 | 28,626,282 | 28,264,944 | 3.56 | 98.03 |
| Salt stress | N_salt01 | 40,034,734 | 39,581,252 | 5.83 | 96.21 |
|  | N_salt02 | 44,533,952 | 44,056,944 | 6.48 | 96.14 |
|  | N_salt03 | 44,718,238 | 44,227,004 | 6.50 | 96.21 |
|  | H_salt01 | 40,847,898 | 40,374,444 | 5.92 | 96.17 |
|  | H_salt02 | 39,929,882 | 39,491,552 | 5.80 | 96.33 |
|  | H_salt03 | 39,319,852 | 38,959,456 | 5.71 | 96.46 |
|  | L_salt01 | 40,036,290 | 39,619,590 | 5.82 | 96.26 |
|  | L_salt02 | 43,389,812 | 42,822,802 | 6.31 | 95.91 |
|  | L_salt03 | 49,465,626 | 48,917,228 | 7.22 | 96.27 |

**Table S6. Summary of clean reads mapping**

| Sample type | Sample name | Total reads | Total mapped | Percentage | Unique mapped | Percentage | Multiple mapped | Percentage |
| --- | --- | --- | --- | --- | --- | --- | --- | --- |
| Different development stages | Egg | 42,881,284 | 24,187,623 | 56.41% | 22,952,326 | 53.53% | 1,235,297 | 2.88% |
|  | Four cells | 42,600,768 | 20,200,370 | 47.42% | 19,278,999 | 45.26% | 921,371 | 2.16% |
|  | Blastulae | 46,718,170 | 25,396,271 | 54.36% | 24,271,023 | 51.95% | 1,125,248 | 2.41% |
|  | Gastrulae | 42,881,284 | 24,187,626 | 56.41% | 22,952,322 | 53.53% | 1,235,304 | 2.88% |
|  | Trochophore | 42,881,284 | 24,187,619 | 56.41% | 22,952,400 | 53.53% | 1,235,219 | 2.88% |
|  | D-stage larvae | 36,163,274 | 6,791,195 | 18.78% | 6,443,107 | 17.82% | 348,088 | 0.96% |
|  | Umbo larvae | 43,044,208 | 24,011,371 | 55.78% | 22,867,699 | 53.13% | 1,143,672 | 2.66% |
|  | Juvenile | 43,739,942 | 25,315,001 | 57.88% | 23,844,190 | 54.51% | 1,470,811 | 3.36% |
| Different adult tissues | Gill | 52,571,902 | 42,408,674 | 80.67% | 39,932,265 | 75.96% | 2,476,409 | 4.71% |
|  | Foot | 51,774,716 | 43,445,527 | 83.91% | 40,362,144 | 77.96% | 3,083,383 | 5.96% |
|  | Adductor muscle | 45,661,534 | 40,224,430 | 88.09% | 36,734,587 | 80.45% | 3,489,843 | 7.64% |
|  | Digestive gland | 45,624,660 | 36,325,819 | 79.62% | 33,701,705 | 73.87% | 2,624,114 | 5.75% |
|  | Mantle | 48,455,080 | 38,728,162 | 79.93% | 35,884,108 | 74.06% | 2,844,054 | 5.87% |
|  | Siphon | 49,368,016 | 41,079,077 | 83.21% | 38,297,681 | 77.58% | 2,781,396 | 5.63% |
|  | Ovary01 | 34,468,792 | 25,108,177 | 72.84% | 23,716,132 | 68.80% | 1,392,045 | 4.04% |
|  | Ovary02 | 35,741,542 | 25,149,497 | 70.36% | 23,736,842 | 66.41% | 1,412,655 | 3.95% |
|  | Ovary03 | 36,258,132 | 25,481,341 | 70.28% | 23,933,412 | 66.01% | 1,547,929 | 4.27% |
|  | Testis01 | 32,024,558 | 21,917,877 | 68.44% | 20,795,814 | 64.94% | 1,122,063 | 3.50% |
|  | Testis02 | 28,336,062 | 20,012,672 | 70.63% | 18,864,576 | 66.57% | 1,148,096 | 4.05% |
|  | Testis03 | 28,264,944 | 19,934,211 | 70.53% | 18,921,720 | 66.94% | 1,012,491 | 3.58% |
| Salt stress | N_salt01 | 39,581,252 | 27,870,270 | 70.41% | 26,456,082 | 66.84% | 1,414,188 | 3.57% |
|  | N_salt02 | 44,056,944 | 29,653,739 | 67.31% | 28,144,306 | 63.88% | 1,509,433 | 3.43% |
|  | N_salt03 | 44,227,004 | 30,710,518 | 69.44% | 29,141,753 | 65.89% | 1,568,765 | 3.55% |
|  | H_salt01 | 40,374,444 | 27,098,825 | 67.12% | 25,794,777 | 63.89% | 1,304,048 | 3.23% |
|  | H_salt02 | 39,491,552 | 27,423,141 | 69.44% | 26,058,023 | 65.98% | 1,365,118 | 3.46% |
|  | H_salt03 | 38,959,456 | 26,790,941 | 68.77% | 25,475,143 | 65.39% | 1,315,798 | 3.38% |
|  | L_salt01 | 39,619,590 | 27,108,102 | 68.42% | 25,736,997 | 64.96% | 1,371,105 | 3.46% |
|  | L_salt02 | 42,822,802 | 29,175,853 | 68.13% | 27,786,168 | 64.89% | 1,389,685 | 3.25% |
|  | L_salt03 | 48,917,228 | 33,534,046 | 68.55% | 31,915,765 | 65.24% | 1,618,281 | 3.31% |

**Table S7. Summary of PacBio full-length transcriptome sequencing**

Please See the Excel file

**Table S8. Summary of the gene prediction results**

| Method | Software | Species | Gene number |
| --- | --- | --- | --- |
| *Ab initio* | Augustus | - | 72,557 |
|  | GlimmerHMM | - | 231,927 |
|  | Genscan | - | 53,981 |
|  | GeneID | - | 35,803 |
|  | SNAP | - | 127,520 |
| Homology-based | GeMoMa | *Branchiostoma floridae* | 39,275 |
|  |  | *Caenorhabditis elegans* | 5,479 |
|  |  | *Crassostrea gigas* | 27,530 |
|  |  | *Ciona intestinalis* | 10,680 |
|  |  | *Drosophila melanogaster* | 5,741 |
|  |  | *Helobdella robusta* | 27,467 |
|  |  | *Homo sapiens* | 12,495 |
|  |  | *Lottia gigantea* | 61,176 |
|  |  | *Octopus bimaculoides* | 18,868 |
|  |  | *Patinopecten yessoensis* | 41,949 |
|  |  | *Strongylocentrotus purpuratus* | 20,929 |
| RNA-seq | Full-length seq | - | 75,225 |
|  | Cufflinks |  | 69,612 |
|  | PASA | - | 30,235 |
| Integration | EVM | - | 40,123 |
|  | PASA-update |  | 26,273 |

**Table S9. Summary of the non-coding RNA annotation**

| Type | Copy | Average length (bp) | Total length (bp) |
| --- | --- | --- | --- |
| miRNA | 968 | 102.75 | 99,462 |
| tRNA | 3354 | 74.58 | 250,141 |
| 18S rRNA | 516 | 321 | 165,636 |
| 28S rRNA | 65 | 115.95 | 7,537 |
| 5S rRNA | 241 | 90.95 | 21,919 |
| CD-box snRNA | 67 | 88.73 | 5,945 |
| HACA-box snRNA | 37 | 183.56 | 6,792 |
| Splicing snRNA | 193 | 147.58 | 28,483 |
